# Covidex: an ultrafast and accurate tool for virus subtyping

**DOI:** 10.1101/2020.08.21.261347

**Authors:** Marco Cacciabue, Pablo Aguilera, María Inés Gismondi, Oscar Taboga

## Abstract

Covidex is an open-source, alignment-free machine learning subtyping tool for viral species. It is a shiny app that allows a fast and accurate classification in pre-defined clusters for SARS-CoV-2 and FMDV genome sequences. The user can also build its own classification models with the Covidex model generator.

**Availability:** Covidex is open-source, cross-platform compatible, and is available under the terms of the GNU General Public License v3 (http://www.gnu.org/licenses/gpl.txt). Covidex is available via SourceForge https://sourceforge.net/projects/covidex or the web application https://cacciabue.shinyapps.io/shiny2/

## INTRODUCTION

There are several tools that have been developed for viral sequence classification, including recently released applications for subtyping of SARS-CoV-2 sequences (Solis-Reyes *et al*., 2018; Randhawa *et al*., 2020; Singer *et al*., 2020; O’Toole *et al*., 2020). Most of them require the alignment of the input data against a set of reference sequences (Altschul *et al*., 1990; Edgar, 2010; Hauser *et al*., 2013). Consequently, these methods can be computationally expensive, particularly for long sequences such as the SARS-CoV-2 genome (>30 Kb). To tackle this problem, alignment-free tools may be used for fast and accurate subtyping (Zielezinski *et al*., 2017).

Current SARS-CoV-2 classification software require the user to be familiar with command line usage, depend on the user to upload the data to an external server or require multiple steps for a correct installation (Singer et al., 2020; O’Toole et al., 2020).Alternatively, Covidex was developed as an open source alignment-free machine learning subtyping tool. It is an app that allows fast and accurate classification of viral genomes in pre-defined clusters. Covidex combines the analytical capacities of the R environment and the user-friendliness of the Shiny user interface (Chang *et al*., 2020). It is available for free and without registration at sourceforge.net for macOS, Linux and Windows users with R and Rstudio dependencies. A portable version is also available for Windows users. Additionally, a web-based version is available at shinyapps.io that can effortlessly be deployed without any installation.

## METHODS AND IMPLEMENTATION

The classification algorithm is divided in three phases. The first phase loads the user data in multi-fasta format and performs the k-mer counting operation using the k-mer package (Figure 1A) (Wilkinson, 2018). If more than 1000 sequences are loaded, this step is performed in parallel by splitting the dataset and processing it in multi-threads to reduce computation time. All tables produced are then merged into one. Each k-mer count is normalized according to the k-mer size and sequence length. In all cases, the k-mer size 6 was selected in order to minimize the computation time (Supplementary Figure 1). The second phase of the classification executes the call to the ranger package predict function using a previously trained random forest model (Wright and Ziegler, 2017). From this, a probability score is calculated as the proportion of trees that supports the corresponding classification. The tool includes a variable threshold to check if each classification meets the desired criteria, by default this threshold is set to 0.6 (for more information see the ROC curves in the Supplementary files 1,2 and 3). The final phase generates a table and a set of statistics plots that are displayed for the user’s ease. The results table is available for download, and snapshots of the plots can also be saved.

**Figure 1.**
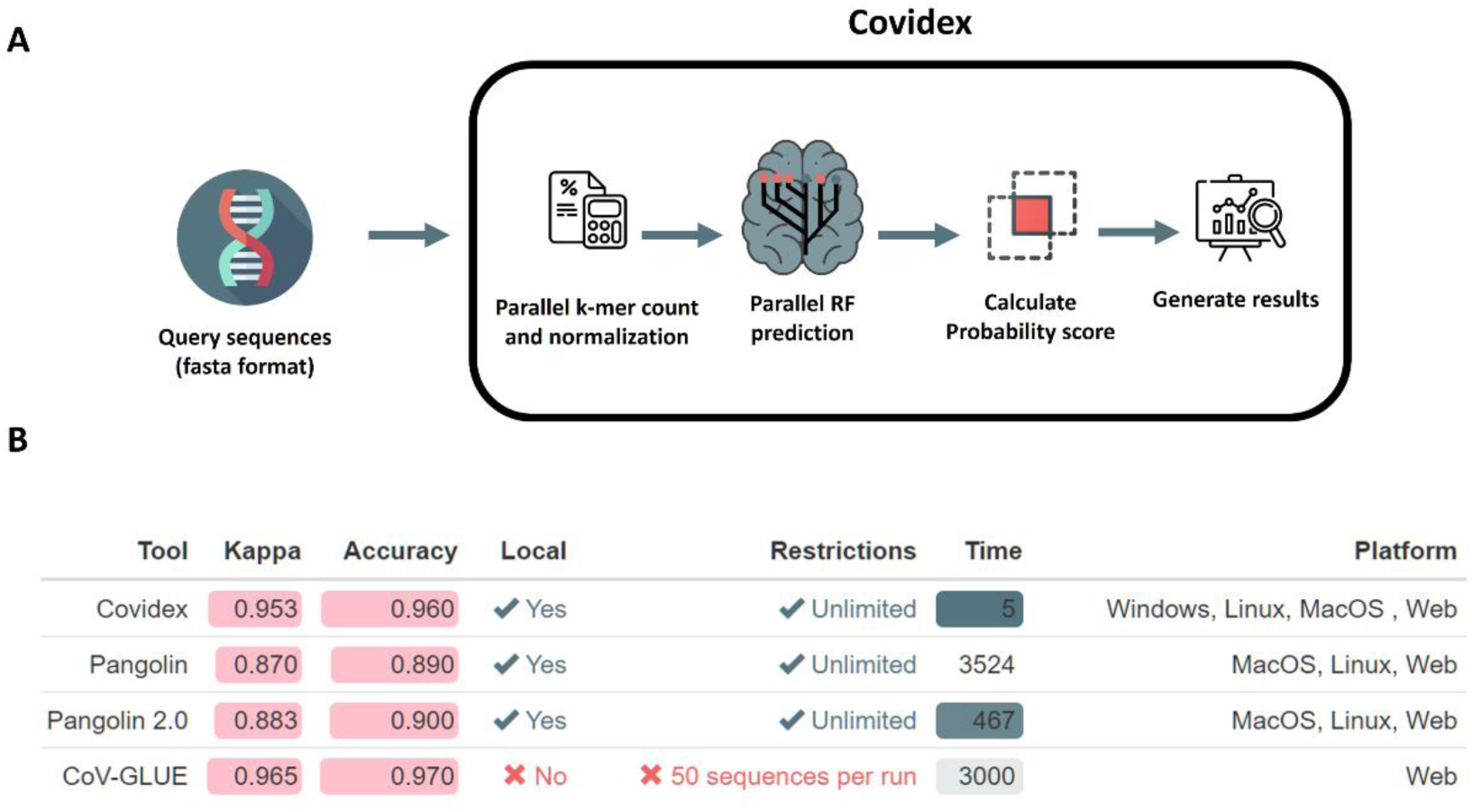
A. Workflow overview of Covidex. First, viral sequences are loaded in fasta format. Next, normalized k-mer counts are obtained from these sequences. The random forest model is then used to classify the query sequences and a probability score based on the number of trees that calls for each class is calculated. Finally, the classification results and plots are generated. B. Comparison of subtyping tools for SARS-CoV-2. Basic statistics were calculated by classifying 100 test sequences with each tool. In the case of CoV-GLUE the sequences was split in two datasets and the application was run sequentially. The calculated statistics were: Cohen’s kappa, as normalized accuracy; accuracy, as total true positive samples over total of samples; and time, in seconds. Local column informs the capability to run each tool in a local host. Icons used were modified from https://www.flaticon.com/authors/freepik

## SOFTWARE DESCRIPTION

Covidex is based on open-source tools and can run on most systems with sufficient memory to run basic java-based applications (Figure 1B). The R shiny environment provides a helpful tool that targets a wide population of scientists with a classification platform that can be easily extended to other viruses. Covidex already includes three classification models that have been developed and tested for SARS-CoV-2 (two) and FMDV virus (one). Previously defined lineage classifications for SARS-CoV-2 and FMDV were used (Hadfield et al., 2018; Rambaut et al., 2020; Brooksby, 1958). In this sense, FMDV and SARS-CoV-2 sequences were downloaded from GISAID and GenBank repositories (Elbe and Buckland-Merrett, 2017; Shu and McCauley, 2017; Clark et al., 2016). For each model, a subset of training sequences and their corresponding class were selected at random retaining 85% of the sequences in each class, providing the class had more than 7 sequences (the R script is available from the authors upon request) (Supplementary Figure 2). Once the random forest model was trained, the remaining 15% of the corresponding sequences of each class was used to test the performance of the trained model (Supplementary files 1, 2 and 3). Alternatively, for classes with less than 7 sequences, all of them were used both in the training and in the test dataset.

The performance of Covidex was compared with two other tools available for SARS-CoV-2 lineage classification: Pangolin and Cov-Glue (Figure 1B). We evaluated the performance of Covidex using a subset of 100 randomly selected sequences from the test dataset classified by Rambaut et al.(Rambaut et al., 2020). Covidex displayed a very high accuracy, similar to that of Cov-Glue (0.96 and 0.97 respectively) and it was the fastest tool, taking only 5 s to complete the task.

To build custom models, Covidex model generator is a companion tool with an intuitive interface that allows users to build their own machine learning classification model in just a few clicks. The user only needs to load a multi-fasta file with known reference sequences and a text file with the corresponding class for each sequence. The tool also allows users to easily change the parameters for building the model (e.g. mtry, number of trees, k-mer size).

## CONCLUSIONS

Covidex combines an accurate and fast machine learning model with an easy-to-use interface (Figure 1B). The computation time of Covidex is extremely low, particularly due to its multi-thread capability, making it a first-choice software for real-time genome subtyping in cases such as the current Covid-19 pandemic. Furthermore, ready-to-use software for analyzing sequences available across all platforms can certainly improve the efficiency of data analyses. Lastly, Covidex’s speed is significantly higher than other classification methods in use. Future improvements include the addition of trained models for other viral species.

## ACKNOWLEDGMENT

We thank Michay Diez for proofreading the article.

