## Supplementary Figure 1 for "Covidex: an ultrafast and accurate tool for virus subtyping"

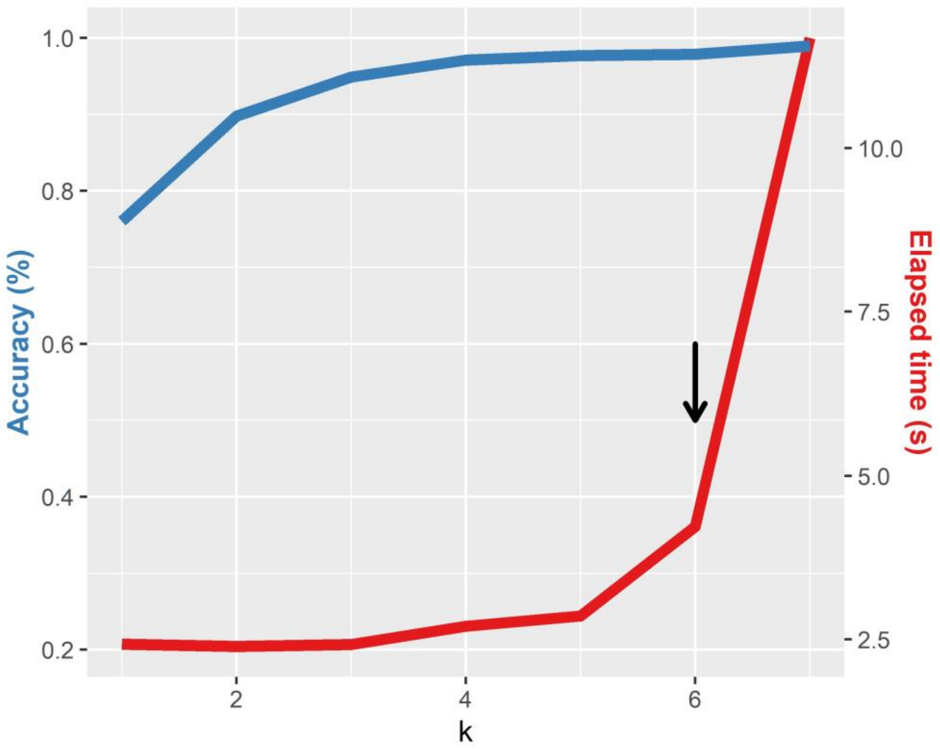


Supplementary Figure 1. Accuracy score and running time for the random forest algorithm, at different values of k, for a set of 1039 whole FMDV genomes from the GenBank database. Black arrow shows the chosen k (highest accuracy with an overall low time).
